# Tannic acid fortifies extracellular matrix against dicarbonyl stress

**DOI:** 10.64898/2026.02.12.705672

**Authors:** Chandrasekar Inbasekar, Claris Niya Varghese, Pavithra Ashok Kumar, Sravanti Uppaluri, Ashok Sekhar, Ramray Bhat

**Affiliations:** Department of Developmental Biology and Genetics, Indian Institute of Science, Bengaluru, 560012, India; Molecular Biophysics Unit, Indian Institute of Science, Bangaluru, 560012 India; Azim Premji University, Bengaluru, 562125, India; Department of Bioengineering, Indian Institute of Science, Bengaluru, 560012, India

**Keywords:** extracellular matrix, methylglyoxal, tannic acid, glycation

## Abstract

Methylglyoxal (MGO), a highly reactive dicarbonyl metabolite that accumulates in diabetes and aging, causes tissue dyshomeostasis, for which therapeutic interventions are limited. Herein, we investigate the potential of tannic acid (TA) in fortifying organ and organismal health against MGO. Anatomical disruption *in vivo* of hydra bodies and *ex vivo* decellularization of murine mesenteries with MGO suggested an impaired interaction between cells and their extracellular matrix (ECM); however, pretreatment of these systems with TA reversed this effect. We confirmed this through subsequent exposure of control and TA-pretreated mammalian cell-secreted endogenous matrix, Collagen I, and basement membrane matrix to MGO. TA prevented loss of ECM biochemical characteristics and restored perturbed cell adhesion and spreading on these substrata induced by MGO. NMR titrations confirmed TA-bound MGO in a 1:5 stoichiometry, potentially quenching its electrophilic properties. Our study posits TA as a novel candidate for protecting organ and organismal architectures against the histopathological effects of dicarbonyl stress.

---

Dicarbonyls, reactive electrophiles formed as intermediates of glucose metabolism, accumulate in blood and various tissues during ageing and diabetes^1^. Methylglyoxal (MGO), one of the most toxic dicarbonyls, reacts with protein and DNA to form cyclic hydroimidazolones and deoxyguanosine adducts, irreversibly altering their structure^2,3^. The detoxification of MGO into D-Lactate is tightly regulated by a highly conserved set of glyoxalase enzymes, GLO 1 and GLO 2^4,5^. Aging and mutation impair glyoxalase activity, which results in impaired detoxification and increased accumulation of MGO, causing excessive crosslinking and dysfunction of cellular biomolecules^6^. Imidazolones are the most common adducts of MGO-protein crosslinking and are collectively called advanced glycated end products (AGE)^7^. AGE accumulation causes increased cellular oxidative stress, drives insulin resistance in diabetes, and even promotes a pro-tumorigenic microenvironment^8^.

Tissues and organs are largely composed of extracellular matrix (ECM), a network of fibrillar and non-fibrillar proteins that provides structural support and preserves tissue specificity and homeostasis^9,10^. Epithelial organs consist of parenchymal cells that localize on a specialized ECM superstructure of Collagen IV, laminin, and proteoglycans known as the basement membrane, and connective tissue cells embedded within a fibrillar collagen matrix^11,12^. ECM remodeling: synthesis, proteolysis, and crosslinking are regulated through cues from the tissue microenvironment^13^. MGO accumulation drives extensive remodeling of the ECM, particularly under metabolic stress, thereby altering the cellular microenvironment by disrupting normal cell-matrix interactions^14,15^. Higher MGO levels exacerbate collagen crosslinking and weaken the basement membrane, aggravating tumor intravasation^16^. The pathological effects of carbonyl stress are manifest specifically because, unlike other posttranslational modifications, glycation does not entail the deployment of conventional enzymes that write or erase their adducts. As a result, cellular strategies such as strengthening the enzyme glyoxalase, which converts MGO to D-lactate through trans-resveratrol and hesperetin, have been under active investigation. However, GLO 1 being an intracellular enzyme, such a strategy will be inadequate for glycation of the ECM; moreover, the expression of the enzyme itself declines with ageing. Therefore, it is crucial to identify inhibitors that effectively neutralize MGO and restore homeostasis within the ECM.

Polyphenols are plant-derived metabolites and well-known free radical scavengers that protect biomolecules against oxidative stress, thereby maintaining cellular homeostasis^17^. Flavonoids such as epigallocatechin gallate (EGCG) and gallic acid are widely used in chronic wound healing because of their strong antioxidant effects^18,19^. Tannic acid (TA), a flavonoid, has been extensively utilized in the food industry as a flavoring agent and in pharmaceutical applications as a drug stabilizer^20^. Owing to its anti-inflammatory properties, it is used as an anti-diarrheal nutraceutical^21^. Moreover, TA is also known to stabilize collagen by strong intermolecular crosslinking, thereby protecting it against proteolytic degradation^22^. However, the pharmacological effects of TA as a putative protectant against the matri-toxic potential of dicarbonyl stress remain unexplored; if established, such observations may open a new window into nutraceutical fortification of organ health through strengthening ECM structure and function. Our results, obtained through investigations using *in vivo*, *ex vivo*, *in cellulo*, and *in vitro* systems, rigorously establish TA as an efficacious dicarbonyl quencher that fortifies the ECM and therefore multicellular organisms and their organs from dicarbonyl-driven dyshomeostasis.

## Results

### TA mitigates MGO-driven disruption of Cell-ECM adhesion

The intracellular physiological concentrations of MGO have been reported to range from 1 to 10 µM^23^. However, under pathological conditions, increased exogenous MGO and impaired detoxification increase its levels to 400 µM^24^. We began our investigations of putative dicarbonyl quenching properties of TA in an *in vivo* model, where dicarbonyl stress has never been explored: hydra, whose body plan, an exemplar for multicellular morphogenesis, consists of a tubular body column between a basal peduncle and an apical region with defined apical tentacles (Figure 1A). Exposing hydra to cadmium chloride (3 µM), a known cytotoxic agent, leads to the dissociation of its body into a slurry of single cells, thus allowing the salt to act as a positive control for the loss of anatomical viability^25^ (Figure 1Bi: stereotypical anatomy of untreated control hydra with a basal stalk, body column, and apical tentacles; 1Bii: hydra body disintegration due to cadmium chloride treatment). We assayed the viability of hydra exposed to progressively increasing MGO concentrations for 4 days (Figure S1) by measuring the proportion of intact live hydra upon chemical exposure. The mean LD_50_ (half lethal dose) of MGO was 1650 μM (Figure 1C). Progressively greater disintegration of hydra was observed upon exposure to 1500 μM (Figure 1Di: stubbed tentacles with body columns) and 1700 μM MGO (Figure 1Dii: absence of tentacles and basal disc with dissociating body column).

**Figure 1:**
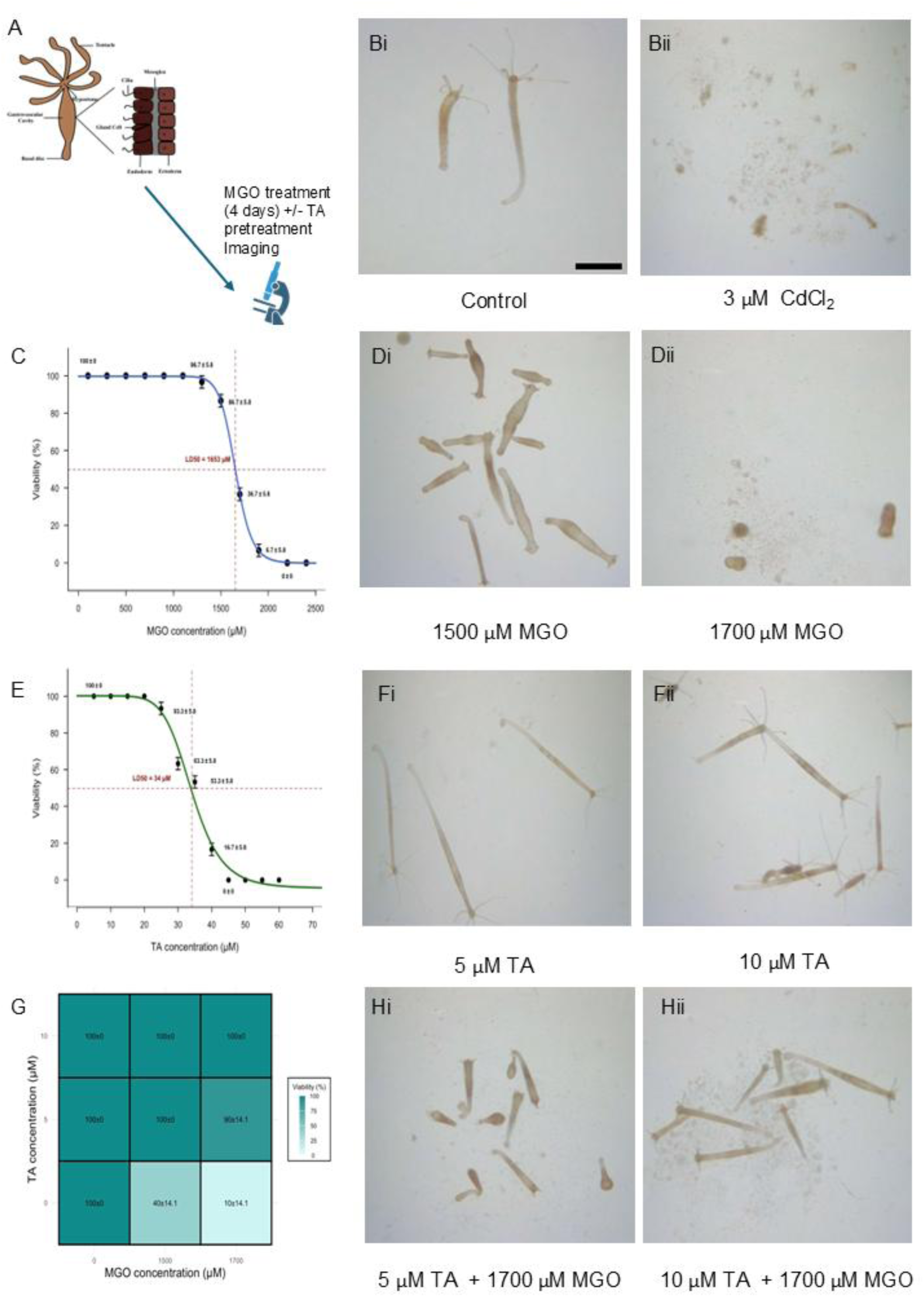
TA rescues disruption by MGO of hydra morphogenesis. (A) schematic depiction of the experimental design for MGO and TA interaction studies in vivo, (B) Bright field micrographs of untreated control hydra (i) and upon treatment with CdCl2 (ii), C) Graph depicting the death curve of hydra upon treatment with MGO, (D) Bright field micrographs hydra upon exposure to 1500 μM (i) and 1700 μM MGO (ii), (E) Graph depicting the death curve of hydra upon treatment with TA, (F) Bright field micrographs hydra upon exposure to 5 μM (i) and 10 μM TA (ii), (G) heatmap depicting the relative mortality of hydra pretreated with 5 and 10 μM TA upon exposure to 1500 and 1700 μM MGO, (H) Bright field micrographs of hydra pretreated with 5 μM TA (i) and 10 μM TA (ii) followed by exposure to 1700 μM MGO, Scale bar for (B, D, F, H) = 1000 μm.

We next measured the effects of TA on hydra viability over 4 days (Figure S2). The mean LD_50_ on the 4th day exposure of the polyphenol was observed to be 35 μM (Figure 1E). Within the first day of exposure, even 30 μM TA did not significantly affect viability, and by the second day, a minor decrease in viability was seen for exposures of 25-30 μM TA. We next tested if the sublethal dosages of TA (5 μM and 10 μM) could protect hydra against MGO toxicity (Figures 1Fi and Fii: intact hydra treated with 5 μM and 10 μM TA). Pretreatment with 5 μM TA significantly reduced MGO-induced mortality, maintaining hydra viability at 100% and 90% at 1500 μM and 1700 μM MGO, respectively; with 10 μM TA, hydra demonstrated complete protection against MGO, preserving normal morphology (Figure 1G: heatmap depicting the relative mortality of hydra pretreated with 5 and 10 μM TA upon exposure to 1500 and 1700 μM MGO; Figures 1Hi and Hii; despite single cells being evident upon treatment with 1700 μM MGO, the hydra bodies seem intact with visible tentacles; compare Hii with Dii, see also Figure S3).

The hydra body plan consists of an ECM called mesoglea, which serves as an anchoring substratum for the organism’s cellular niches (the endo- and ecto-derms)^26^; the dissolution of the body under exposure to MGO suggested its deleterious effect on cell-ECM adhesion. We tested this hypothesis further within murine mesenteries: ECM-rich mesothelia-covered organs that are part of the vertebrate peritoneum^27^. Mesenteric explants were dissected from 4-6 week-old BALB/c mice and cultured *ex vivo* (Figure 2A). We optimized the concentration of MGO to 200 μM based on a concentration-diverse screen: treated mesenteries exhibited significant mesothelial shedding (Figure 2Bi-ii). Mesothelial confluence in samples treated with 10 µM TA, with or without subsequent MGO exposure, remained comparable to untreated controls, indicating that TA effectively prevented the MGO-induced loss of mesothelial adhesion on mesenteries (Figure 2Biii-iv and Figure 2C). Taken together, the pharmacological assays with both our *in vivo* and *ex vivo* systems indicated an MGO-driven attenuation of cell-ECM adhesion that was prevented by a pretreatment with TA.

**Figure 2:**
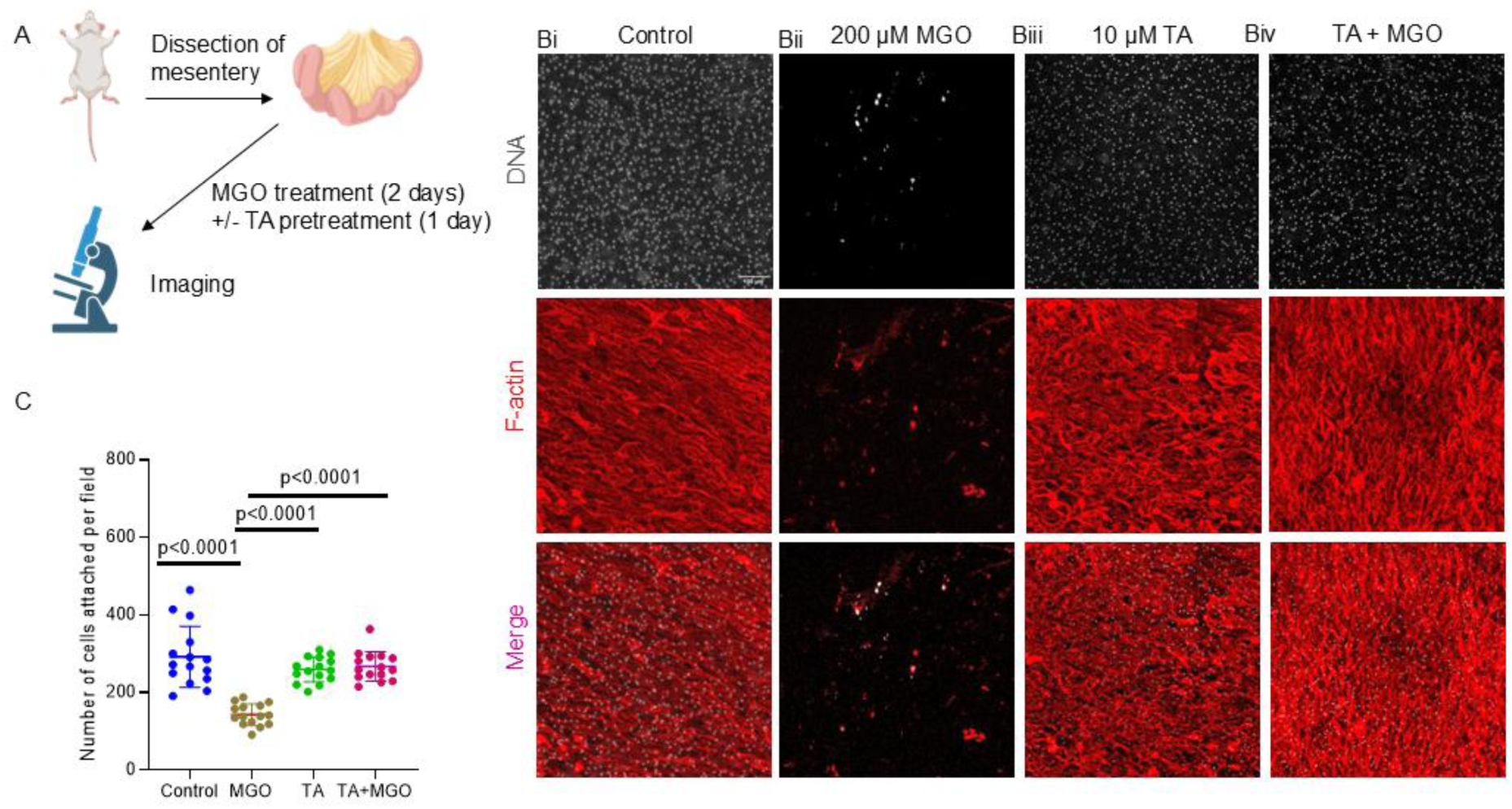
TA rescues disruption by MGO of cell-ECM adhesion. (A) Schematic representation of the isolation of murine mesenteries and subsequent treatment with MGO and TA, (B) Spinning disc confocal micrograph of murine mesenteric tissue: (i) untreated control, (ii) treated with 200 µM MGO for 4 days, (iii) treated with 10 µM TA for 1 day, and (iv) exposed to MGO after TA treatment; mesothelia stained with DAPI for DNA (white) and with Phalloidin for F-actin (red), (C) a graph representing number of cells attached per field upon the above treatments; p < 0.0001 for all; significance computed using one way ANOVA with Tukey’s post hoc comparison; error bars represent mean ± SD. Scale bar for B: 100 μm, experiments repeated n=3 or more times.

### TA protects Cell-Secreted Endogenous Extracted Matrix (CEEM) from MGO-induced disruption

To mechanistically dissect how MGO regulates cell-matrix interactions, we performed assays with untransformed human cell-secreted endogenous extracellular matrices (CEEM). Immortalized untransformed human fallopian tubal epithelial cells (hTERT-FT) were cultured till confluence, osmotically decellularized using 20 mM ammonium hydroxide to obtain CEEM, visualized through crystal violet staining (Figure 3A). Untreated CEEM exhibited a mesh-like pattern, which was damaged and fragmented upon exposure to 200 µM MGO for 4 days. 100 µM TA pretreatment for 1 day, followed by with and without subsequent MGO exposure, showed little fragmentation, suggesting that TA effectively prevents MGO-induced matrix degradation (Figure S4). The protein concentration of the CEEM recorded through absorbance of crystal violet in the washout samples showed a significant decrease upon MGO treatment that was restored to near control values upon pretreatment of CEEM with TA (Figure S5). Our observations suggested that MGO may induce a degradative effect on the CEEM that was thwarted by TA pretreatment.

**Figure 3:**
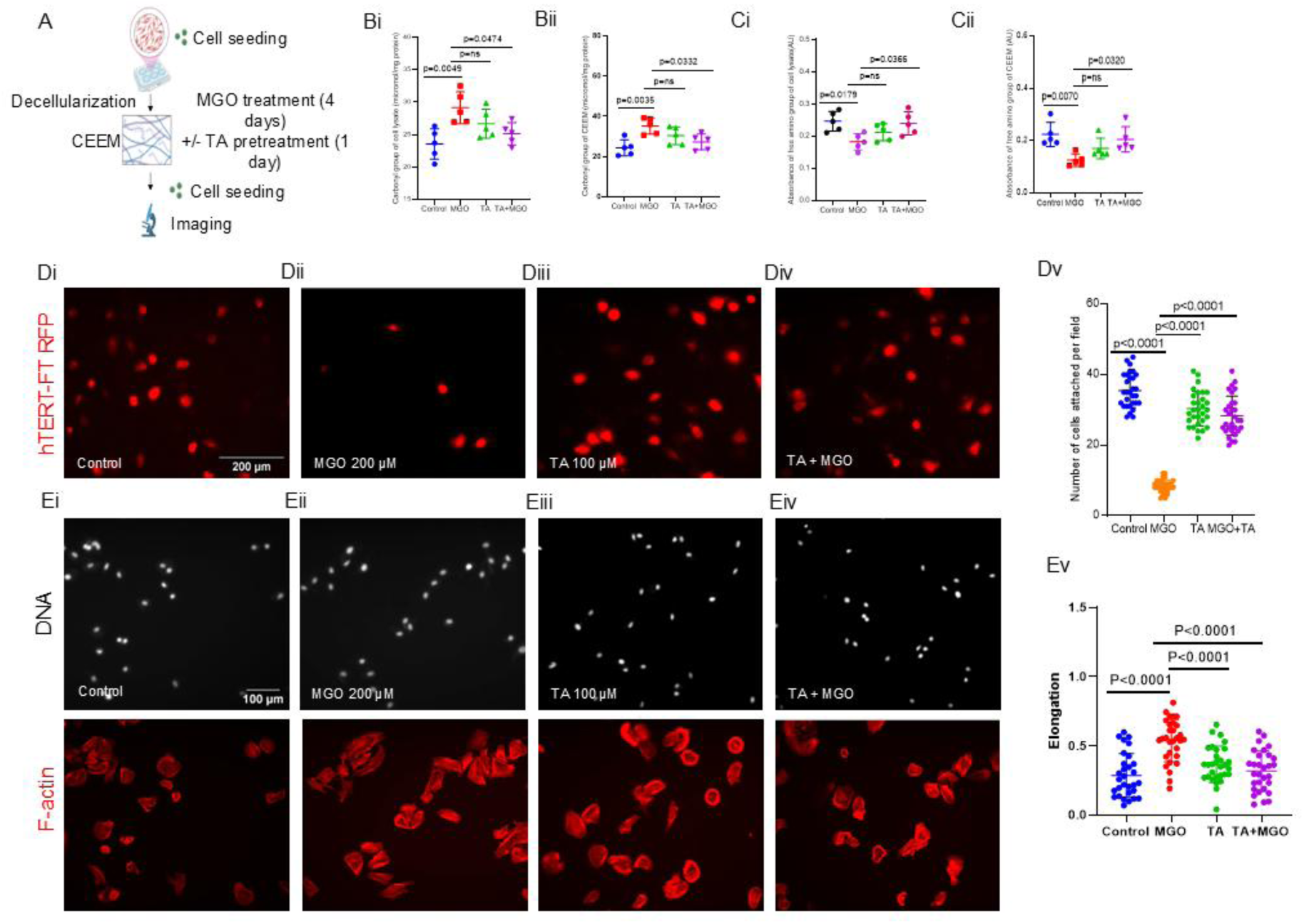
TA rescues disruption by MGO of the adhesion of cells to human cell-secreted ECM. (A) Schematic representation for the preparation of cell-extracted matrix (CEEM) for MGO and TA interaction studies, (Bi) Carbonyl content of hTERT-FT cells (Bii) Carbonyl content of CEEM, control, treated with 200 µM MGO for 4 days, treated with 100 µM TA for 1 day, and exposed to MGO after TA treatment. (Ci) Free amine content of hTERT-FT cells (Cii) Free amine content of CEEM, control, treated with 200 µM MGO for 4 days, treated with 100 µM TA for 1 day, and exposed to MGO after TA treatment (D) represents spinning disc confocal micrographs showing adhesion of RFP labelled hTERT-FT cells on CEEMs: (i) control, (ii) treated with 200 µM MGO for 4 days, (iii) treated with 100 µM TA for 1 day, and (iv) exposed to MGO after TA treatment, (v) represents a graph showing mean cell number per field measured upon the different chemical treatments in Di-iv; (E) spinning disc confocal micrographs showing spreading of hTERT-FT on CEEM: (i) control, (ii) treated with 200 µM MGO for 4 days, (iii) treated with 100 µM TA for 1 day, and (iv) exposed to MGO after TA treatment, cells stained with DAPI for DNA (white) and with Phalloidin for F-actin (red), (v) graph for elongation of hTERT-FT cells on CEEM under treatments in Ei-iv. Significance for all graphs computed using one-way ANOVA with Tukey’s post hoc pairwise comparisons; error bars represent mean ± SD. Scale bar for D: 200 μm and for E:100 μm, experiments repeated n = 3 or more times.

Next, we studied the biochemical alteration induced by MGO in hTERT-FT cells and CEEM. We observed that both cell lysate and CEEM exposed to MGO showed significantly higher carbonyl levels (assessed through conversion of dinitrophenylhydrazine to dinitrophenylhydrazone upon binding aldehyde/ketone groups), whereas subsequent MGO exposure to TA-pretreated cell lysate and CEEM exhibited carbonyl levels comparable to control, suggesting that TA effectively inhibits MGO-induced adduct formation in the lysates of cells and CEEMs (Figures 3Bi-ii). Free amine levels in lysates of cells and CEEM (assessed through their binding to p-benzohydroquinone) showed a pattern that was opposite to carbonyl levels (Figures 3Ci-ii).

The MGO-induced restoration of substrate-free amines in CEEM prompted us to investigate untransformed cell adhesion on MGO-treated CEEMs, with or without TA pretreatment. 200 µM MGO exposure for 4 days reduced substrate adhesion of hTERT-FT cells; treatment with TA with and without subsequent MGO exposure restored cell-ECM adhesion (Figure 3Di-iv and Figure 3Dv). Similar results were also obtained with the untransformed human peritoneal mesothelial line MeT-5A (Figure S6).

Untransformed epithelial cells spread on native extracellular matrices through reinforcement of basoapical polarity using hemidesmosomal adhesion, with greater contact area on substrata that exhibit cognate extracellular ligands and/or exhibit high levels of stiffness^28^. Therefore, we assayed for the relative contact area (elongation) of hTERT-FT cells cultured on CEEM. Cells on the MGO-treated CEEM (200 µM; 4 days) exhibited greater spreading than control cells (Figure 3Ei-ii). In contrast, cells on TA-treated CEEM (100 µM; 1 day) with or without subsequent MGO exposure maintained a predominantly circular morphology with spread that was closer to control cells (Figure 3Eiii-iv and Figure 3Ev). Similar results were observed for another immortalised human peritoneal mesothelial line, MeT-5A (Figure S7).

### TA protects Cell-Matrix interactions on Collagen I and LrBM from MGO-induced disruption

The alteration in cell-ECM interactions on native matrices is difficult to interpret, given that such matrices are composed of several proteins, each with its own capacity to regulate cellular traits. To further elucidate the molecular mechanism underlying these observations, we investigated how MGO alters cellular interactions with specific ECM proteins that principally constitute the tissue extracellular milieu and have been well characterized for their regulation of cell adhesion and migration, i.e., on MGO-treated and TA-treated Collagen I and laminin-rich basement membrane matrices (Figure 4A).

**Figure 4:**
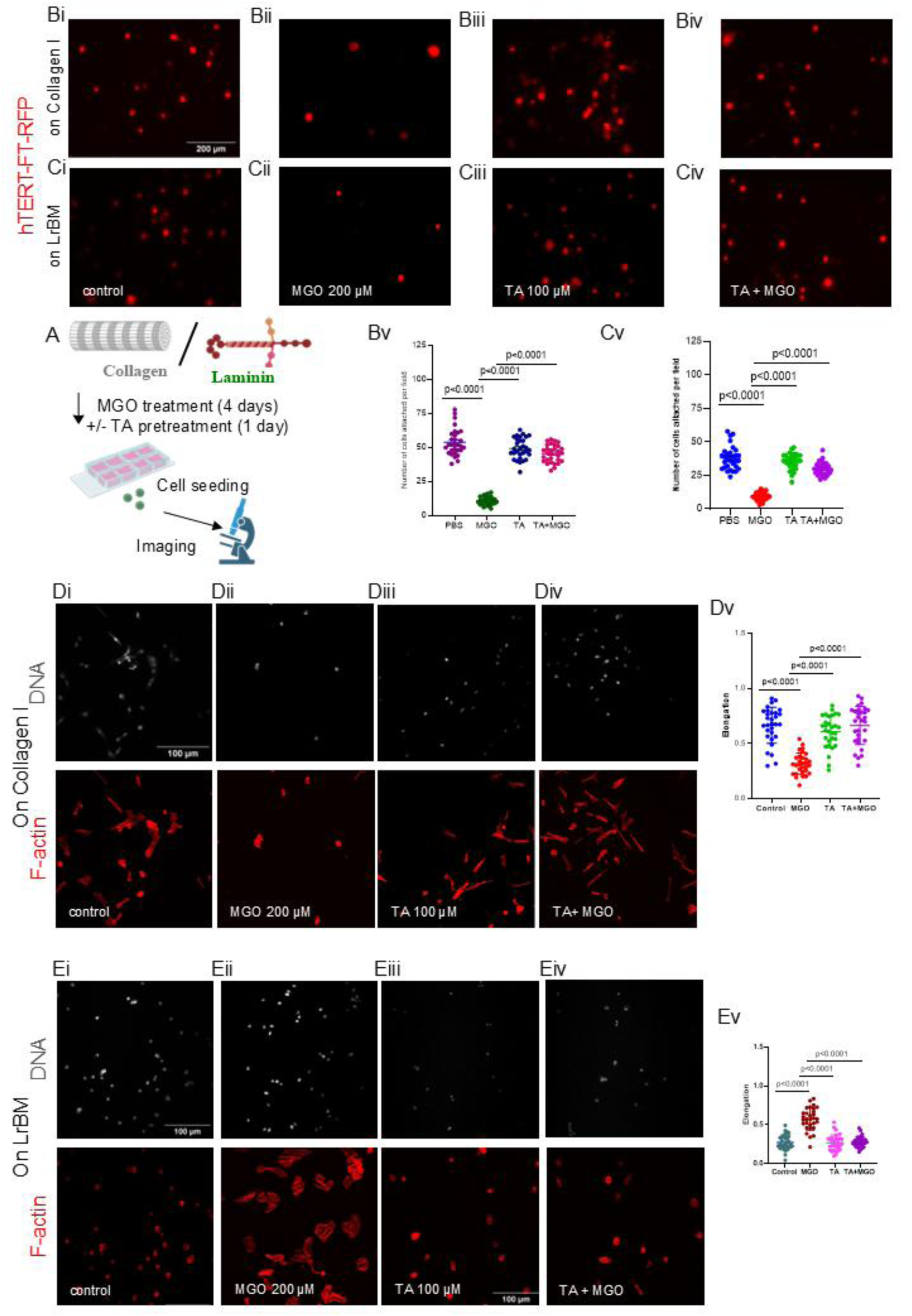
TA rescues disruption by MGO of cell adhesion to Collagen I and LrBM. (A) Schematic representation for the LrBM/Collagen I gel preparation for cell adhesion, (B) and (C) represent spinning disc confocal micrographs showing adhesion of RFP labeled hTERT-FT on Collagen I and LrBM gels, respectively: (i) control, (ii) 200 µM of MGO for 4 days, (iii) treated with 100 µM of TA for 1 day, and (iv) MGO exposure with TA pretreatment. Bv and Cv represent graphs showing the mean cell number per field measured for the different chemical treatments. (D) represents spinning disc confocal micrographs showing spreading of hTERT-FT on Collagen I: (i) control, (ii) 200 µM of MGO for 4 days, (iii) treated with 100 µM of TA for 1 day, and (iv) MGO exposure with TA pretreatment; cells stained with DAPI for DNA (white) and with Phalloidin for F-actin (red). (Dv) represents graphs for elongation of hTERT-FT cells on Collagen I gels under different treatments. (E) represents spinning disc confocal micrographs showing spreading of hTERT-FT on LrBM gels: (i) control, (ii) 200 µM of MGO for 4 days (iii) treated with 100 µM of TA for 1 day and (iv) MGO exposure with TA pretreatment; cells stained with DAPI for DNA (white) and with Phalloidin for F-actin (red), (Ev) represent graph for elongation of hTERT-FT cells on LrBM gels under various treatments; significance for all graphs computed using one-way ANOVA with Tukey’s post hoc pairwise comparisons; error bars represent mean ± SD. Scale bar for B, C: 200 μm and for D, E: 100 μm, experiments repeated n = 3 or more times.

hTERT-FT-RFP and MeT-5A-RFP cells were added to 1 mg/mL polymerized Collagen I and 8-10 mg/mL polymerized LrBM substrata, and their relative adhesion was measured using fluorescence. For both cell types, adhesion was reduced upon MGO treatment (200 µM; 4 days). TA pretreatment normalized cell adhesion on the substrate, aligning it with control levels (Figures 4Bi-v and 4Ci-v: adhesion of hTERT-FT cells on Collagen I and LrBM gels). The same trends were observed in Met-5A cells (see Figures S8i-ii and S9i-ii).

To study whether glycation affected cell spreading, hTERT-FT and MeT-5A cells were seeded on Collagen I gels, and their contact area with their substratum was estimated after 6 h. On MGO-treated gels, the morphology of both hTERT-FT and MeT-5A appeared rounded and exhibited reduced spreading relative to the untreated control, suggesting depleted binding sites due to glycation by MGO. Cells on substrates treated with TA, with or without subsequent MGO exposure, exhibited contact areas similar to those of untreated controls (Figure 4Di-v and S10i-v).

hTERT-FT and MeT-5A cells cultivated on control LrBM matrix exhibited a rounded, less spread morphology due to a relatively greater hydrophobicity on such surface milieu compared with Collagen I, whose acidic amino acids support cell spreading. Cells showed greater spreading on MGO-treated LrBM gels, which could be explained by a mechanical alteration of the scaffold or due to dissolution of the LrBM matrix^16^, which was prevented with TA pretreatment (Figure 4E and S11: spreading of hTERT-FT and MeT-5A on Collagen I gels: control (Ei, S11i), MGO (Eii, S11ii), TA treatment (Eiii, S11iii), and MGO with TA pretreatment (Eiv, S11iv) cells stained with DAPI for DNA (white) and with Phalloidin for F-actin (red)); Figure 4Ev and S11v: graphs for elongation of hTERT-FT and MeT-5A cells on LrBM matrix).

### TA rescues the alteration by MGO of the matrix structure

The impaired adhesion and spreading of epithelial and mesothelial cells on MGO-treated Collagen I and LrBM matrices suggested an alteration in the molecular functions of these ECM proteins by glycation, thereby reducing the integrin binding site. To directly assess these structural changes, acellular Collagen I gels were subjected to MGO and TA pretreatments and stained with a collagen-binding dye: picrosirius red (Figure 5A). Decreased intensity of picrosirius red was observed in Collagen I gel treated with MGO, in comparison to TA pretreated gels (Figure 5B and C). To quantify the extent of degradation, levels of Collagen I that leached from the gels were measured. MGO treatment caused maximal Collagen I depletion, likely resulting from enhanced matrix degradation, an effect effectively blocked by TA pretreatment. This confirms that TA-stabilized collagen was less susceptible to degradation (Figure 5D).

**Figure 5:**
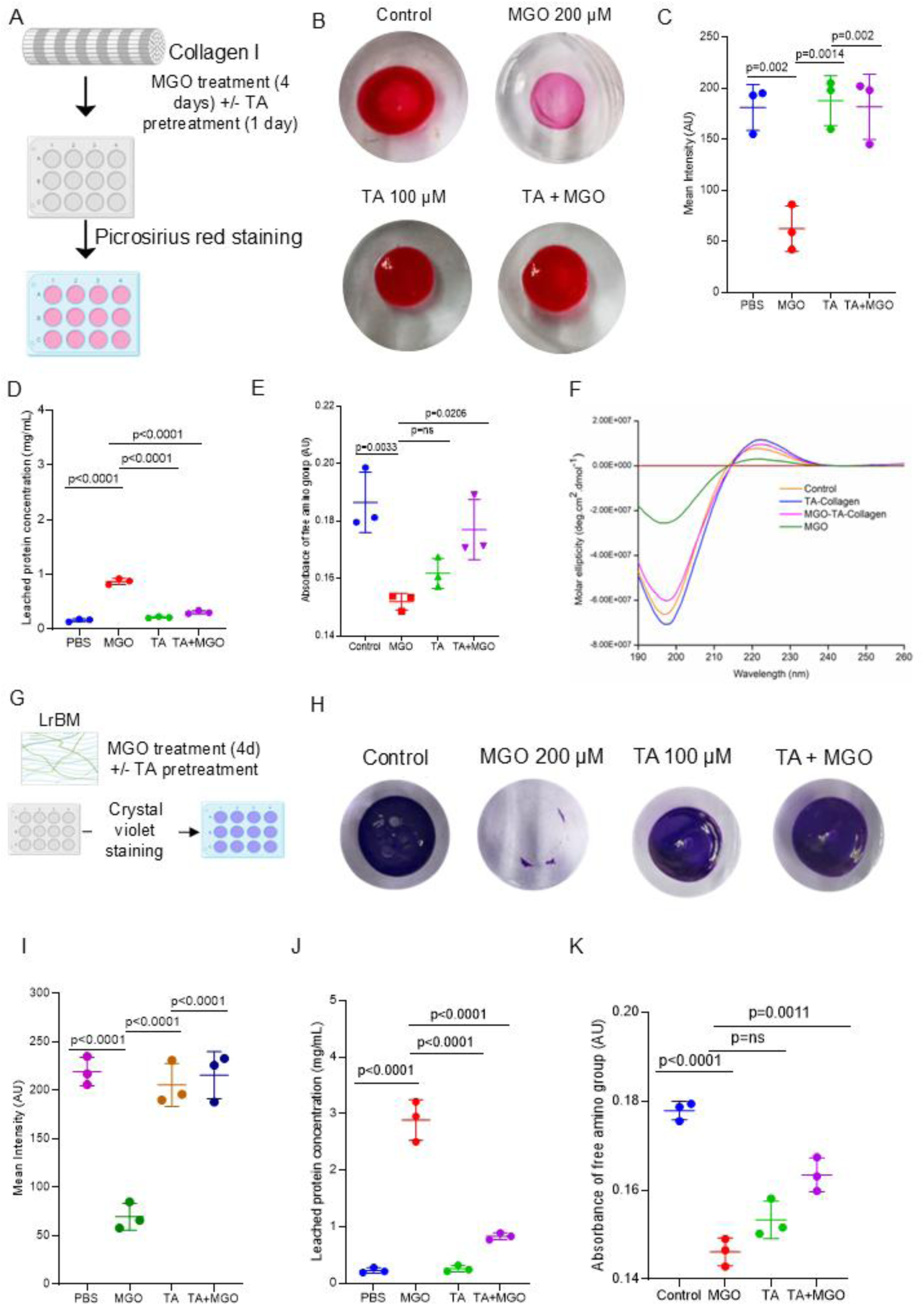
TA restores alterations by the MGO of the matrix structure. (A) Schematic depiction for the Collagen I gel preparation for MGO and TA interaction studies (B) Digital images of polymerized Collagen I gel, control (top left), 200 µm of MGO for 4 days (top right), 100 µm of TA for 1 day (bottom left), 100 µm of TA for 1 day and 200 µm of MGO for 4 days (bottom right), stained with picrosirius (red), (C) Graph for mean intensity of picrosirius staining of Collagen I gels stained in (B), (D) Graph showing mean levels of leached protein from Collagen I gel under treatments in (B), (E) Graph showing mean absorbance of free amine groups of Collagen gels under treatments in (B), (F) CD spectra of Collagen I under various treatments in (B), (G) Schematic depiction of LrBM gel preparation for MGO and TA interaction studies, (H) Digital images of solidified LrBM gels, control, treated with 200 µm of MGO for 4 days, with 100 µm of TA for 1 day, and with 100 µm of TA for 1 day and 200 µm of MGO for 4 days (from left to right), stained with crystal violet (blue), (I) Graph for mean intensity of crystal violet staining of LrBM gels stained in (H), (J) Graph showing mean levels of leached protein from LrBM gel under treatments in (H), (K) Graph showing mean absorbance of free amine groups of LrBM gels under treatments in (H), One way ANOVA Tukey’s multiple comparison was performed for all graphs. Error bars represent mean ± SD. Experiments performed n = 3 times.

MGO-treated gels exhibited reduced free amine absorbance, validating their tendency to crosslink ECM arginine and lysine residues. TA pretreatment restored free-amine absorbance, in accordance with the picrosirius red staining results (Figure 5E). We next investigated the effect of MGO and TA on the secondary structure of Collagen I using circular dichroism (CD) spectroscopy. Monomeric Collagen I (pH 4.7) exhibited two distinct peaks, a positive peak at 222 nm and a negative peak at 197 nm, of polyproline II helical conformation^29^. MGO treatment significantly decreased both peak intensities, indicating an alteration of the triple helical conformation and consequent structural alteration. In contrast, we observed insignificant changes in the peak of samples incubated with TA with or without subsequent MGO exposure, suggesting that TA preserves and stabilizes Collagen structure, even in the presence of MGO (Figure 5F).

Unlike Collagen I, which self-assembles into high-order crosslinked fibrillar structures, LrBM forms a soft and non-fibrillar gel, primarily composed of laminins and Collagen IV, and is stabilized predominantly through weak non-covalent interactions^30^. To assess stability upon MGO treatment, LrBM gels were stained with crystal violet (Figure 5G). MGO treatment led to the complete dissolution of LrBM, as seen by the absence of crystal violet stain (Figure 5H). Consistent with its protective role on Collagen I, TA pretreatment stabilized the LrBM matrix and prevented MGO-induced degradation (Figure 5I). We quantified the concentration of proteins leached from the treated gels. MGO released the highest level of proteins from the gels, likely due to glycation-induced ECM disruption, an effect that was reduced by TA pretreatment, highlighting the matrix-protective capacity of TA (Figure 5J). In line with its reactivity toward Collagen I, MGO significantly reduced free-amine absorbance in LrBM, reflecting its interaction with the matrix. However, TA pretreatment restored higher free amine absorbance (Figure 5K).

### NMR titration reveals a direct interaction between TA and MGO

To characterize the direct interaction between MGO and TA, we performed titrations detected using ^1^H 1D NMR spectra. Aiming to probe the MGO peaks in NMR spectrum, we prepared 18 samples containing 333 μM MGO and [TA]/[MGO] concentration ratios ranging from 0 to 5 (Figure 6A). In aqueous solution, MGO exists in equilibrium with its monohydrate and dihydrate forms (Figure 6B)^31^. The ^1^H 1D spectrum of MGO in the absence of TA (Figure 6C) contains peaks from dehydrated MGO (1), MGO monohydrate (2), and MGO dihydrate (3) at distinct chemical shifts, as these three forms are in slow exchange on the NMR timescale. As we increase the [TA]/[MGO], we observe a decrease in the unbound methyl proton peak intensity of all three forms of MGO. This indicates that the MGO is interacting with TA, thereby reducing the amount of MGO in the unbound state. The methyl peaks of MGO monohydrate and dihydrate in the unbound state are isolated from other peaks and can be quantified during the course of titrations (Figure 6D top). Between 0.2 and 0.28 equivalents of TA, the unbound MGO peaks disappear completely, suggesting that all MGO molecules are bound to TA beyond [TA]/[MGO] of 0.28. This implies an approximate stoichiometry of one TA molecule interacting with four to five molecules of MGO.

**Figure 6:**
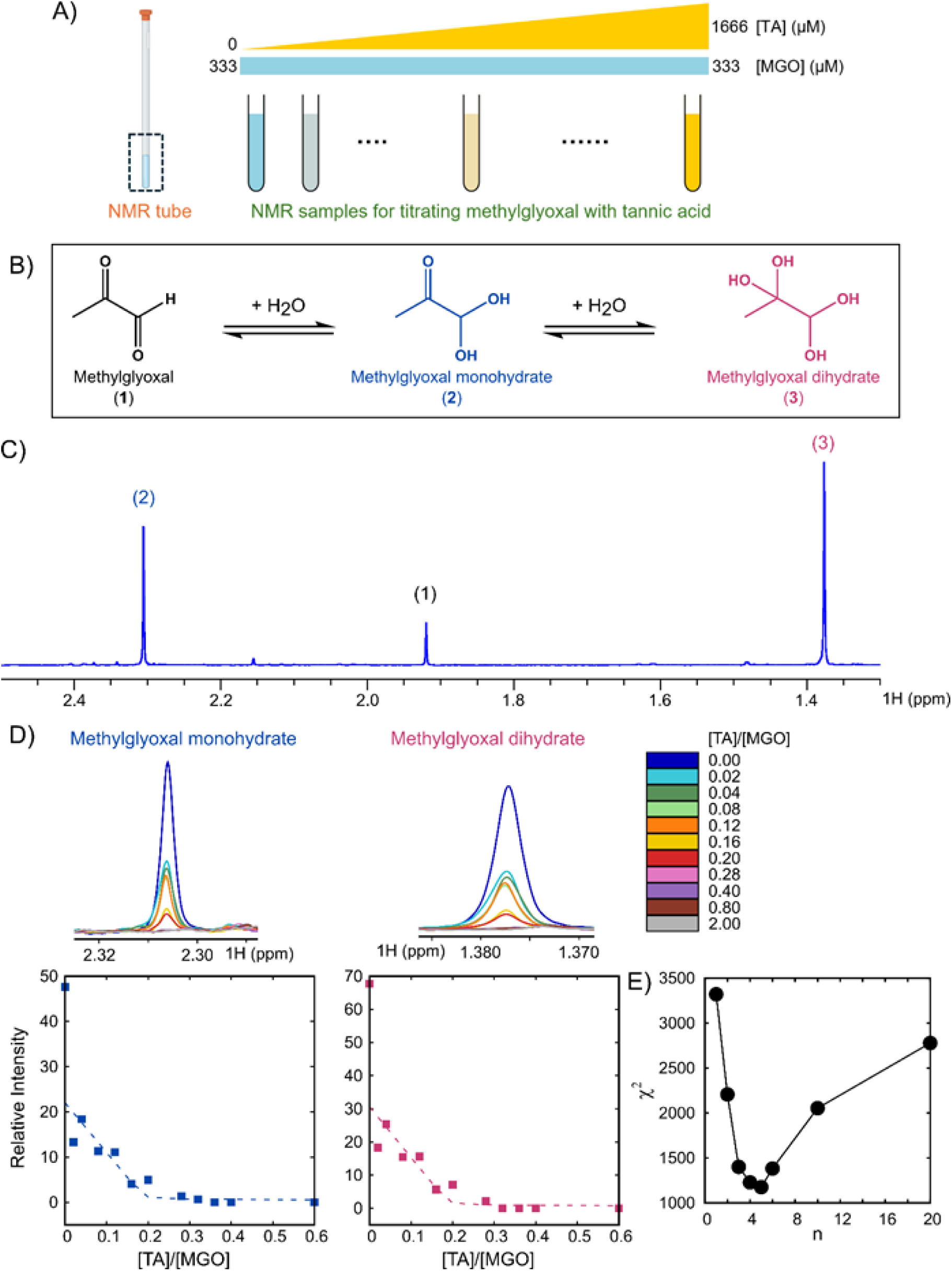
NMR titration reveals a direct interaction between TA and MGO. (A) Schematic demonstrating the sample preparation strategy for NMR titrations. The concentration of the methylglyoxal (MGO) is 333 μM across all NMR samples, while the concentration of the tannic acid (TA) is varied from 0 to 1666 μM across the titration. (B) The hydration reaction of methylglyoxal in aqueous solution. (C) 1H 1D NMR spectrum of MGO showing three distinct peaks for the methyl groups of forms (1), (2), and (3) from panel C. (D) 1H 1D spectrum as a function of TA concentration (top) and titration plot (bottom) of methyl 1H peak of methylglyoxal monohydrate (left) and methylglyoxal dihydrate (right). The dotted line in the titration represents data points fit to a 5-site (1:5 TA:MGO) binding model. (E) χ2 surface for the stoichiometry of interaction (n) using an n-site binding model. The minimum in *χ*^2 occurs at a value of n=5.

NMR titration profiles graphing the decay of the unbound peak intensity are sensitive to stoichiometry (1: n) of the interaction. The titration profiles of methyl resonances from unbound MGO monohydrate and MGO dihydrate, obtained by plotting their relative intensities against [TA]/[MGO] (Figure 6D bottom), show a progressive decay of peak intensity to a final value of 0. We have used an n-site binding model to globally fit the titration profiles of the two hydrated forms of MGO. Although the sharp intensity drop from [TA]/[MGO]=0 to [TA]/[MGO]=0.02 cannot be fit well using this model, the stoichiometry of the TA: MGO complex can be estimated reliably as it depends on the titration point where the unbound peak intensity goes to zero, under conditions where the interaction is strong. The plot of the *χ*^2^ Metric as a function of n shows a clear minimum at n=5, confirming that the stoichiometry of the interaction in the TA: MGO complex is 1:5 (Figure 6E).

## Discussion

Our study was inspired by two landmark efforts in 2024: the first by Kong and coworkers showed how MGO could bring about proteolysis of BRCA2 in cells, eliciting its functional haploinsufficiency, leading to malignant transformation^24^. The second paper, by Fan and coworkers, showed how the accumulation of AGE triggered by the Maillard reaction mediated by MGO with proteins alters collagen architecture and its mechanical response. In combination with oncogenic β catenin signaling, such a tissue microenvironment induced the progression of hepatocellular carcinoma^32^. These papers clearly motivate efforts to develop compounds that may fortify the extracellular matrix against electrophilic attacks by dicarbonyls. Our study positions TA as a strong quencher of the deleterious effects of MGO on all the above-examined spatial morphological scales. In doing so, the experiments in this manuscript also represent a composite and self-sufficient set of assays that can be deployed as high-throughput screens to test the putative effects of natural and synthetic compounds against dicarbonyl stress.

The protective effect of TA against MGO is evident in the *in vivo* Hydra model, whose mesogleal matrix forms a central interstitial layer between the ectodermal and endodermal cell layers and contains type I and type IV Collagen as well as laminin^33^. The expression of ECM proteins has been shown to be critical for hydra morphogenesis^34^. In our study, we observed that exposure to MGO caused extensive disintegration within 24 hours, resulting in nearly 100% organismal death. This effect is likely mediated by the rapid shedding of cells from their ECM substrata, as nearly intact mesoglea were frequently observed floating immediately upon MGO treatment. However, TA pretreatment preserved hydra viability even at MGO concentrations close to the predetermined LD₅₀. Based on the evidence we have gathered, it is likely that MGO interferes with cellularization of the mesoglea in hydra, and similar to the *in vitro* cellular adhesion assays, TA blocks this effect. This was supported by the experiments with murine mesenteries, which, when treated with MGO, exhibited significant cell shedding, likely resulting from glycation-induced conformational changes in their ECM binding proteins. TA pretreated mesenteries, even when exposed to MGO, exhibited higher cell numbers comparable to the untreated control tissues.

Our biochemical assays using cell-secreted endogenous extracellular matrix (CEEM) further confirmed the mechanisms of MGO toxicity. The marked reduction in protein content, elevated carbonyl accumulation, and reduced free amine level in MGO-treated CEEMs (which phenocopy the effects of MGO with LrBM) is consistent with glycation-driven protein carbonylation in diabetes. The electrophilic MGO could potentially bind the nucleophilic basic amino acids of collagen, disallowing the binding of anionic picrosirius to the same protonated amino acids. Therefore, the crosslinked glycated Collagen I is unable to retain the stain of picrosirius, as has also been reported earlier^16^. Integrin α2β1, a transmembrane receptor, facilitates cell adhesion by binding to the arginine-containing domains of ECM proteins such as collagen and laminin^35–37^. Since MGO crosslinking targets arginine primarily, decreasing the number of accessible binding sites for integrin recognition substantially reduces, attenuating cell adhesion. The multiple phenolic groups of TA are capable of forming hydrogen bonds predominantly with the amine group of lysine and arginine residues in proteins, rendering them resistant against glycation. Therefore, TA pre-treatment competitively blocks the arginine sites and prevents the structural changes induced by MGO, which result in higher cell attachment in the pretreated gels. These results were consistent with earlier reports demonstrating a concentration-dependent decrease in the arginine-hydroxyproline ratio in collagen following MGO exposure, reflective of selective arginine modification^38^. LrBM is mainly composed of Laminins and Collagen IV, stabilized predominantly through weak non-covalent interactions. MGO treatment causes dissolution of the LrBM matrix, resulting in poorer initial adhesion of the cells but later higher migration of the adhered ones on the hard plastic substrata^39,40^.

Polyphenols possess hydroxyl-rich aromatic rings that can trap the highly reactive dicarbonyls and convert them into less reactive phenolic adducts^41,42^. NMR titrations reveal that TA interacts with MGO to form a 1:5 TA-MGO complex. TA is a glucose derivative with its hydroxyl groups covalently attached to five digallol (3, 4, 5-trihydroxylbenzene) ester groups. Each of the five terminal gallol moieties in TA can interact with one MGO molecule, resulting in the observed 1:5 stoichiometry of the TA: MGO interaction. Similar to other polyhydroxyl aromatic compounds with pyrogallol (1, 2, 3-trihydroxylbenzene) and gallic acid (3, 4, 5-trihydroxybenzoic acid) moieties, TA likely interacts with MGO through a covalent adduct formation. Pyrogallol reacts with MGO via an electrophilic addition at the meta position to C3, with a second addition either at ortho or at para positions to C3 under neutral or alkaline conditions. However, gallic acid shows weaker reactivity due to its carboxylic acid substitution at the meta position of C3, and still forms one MGO conjugated adduct^43^. In TA, the meta position of the gallol groups is engaged in ester linkages to the glucose core, suggesting that electrophilic addition of MGO occurs at either at ortho or at para position to C3 of the gallol groups^44–46^.

Aminoguanidine (AG), a hydrazine derivative, is a well-known drug for diabetic nephropathy. AG prevents the protein glycation products by trapping reactive dicarbonyls and competitive binding, thereby preventing AGE formation. Despite its therapeutic potential, AG causes severe side effects, including anemia, gastrointestinal infection^47^. We chose tannic acid due to its approved use as a flavoring and clarifying agent in foods. Our investigations in this manuscript demonstrate little cellular or organ toxicity associated with the used concentration of TA; however, detailed histopathological studies will be performed to further confirm this in the future. On the other hand, the ability of TA to be able to bind MGO in a 1:5 ratio shows promise for its efficiency to ameliorate glycative stress caused by the latter.

In summary, our study provides novel mechanistic insight into how dicarbonyl stress contributes to tissue degeneration and highlights TA as a promising natural molecule for preventing MGO-induced ECM degradation. Although our assays provide clear evidence on the effect of MGO-induced glycation on ECM proteins and the protective role of TA, we would like to extend our assays to other dicarbonyls that are known to accumulate in tissues, such as glyoxal and 3-deoxyglucosone. Building on the presented findings, future investigations of glycation-preventing roles of polyphenols could open up a new therapeutic window into the management of diabetic organopathies and end-stage disease.

## Materials and methods

Methylglyoxal, p-benzoquinone, ammonium hydroxide, glutaraldehyde, formaldehyde, and tannic acid were procured from Merck. 2,4-dinitrophenyl hydrazine, urea, and solvents were procured from Sisco Research Laboratories. Trichloroacetic acid, crystal violet, and PBS buffer were procured from HiMedia.

### Cell culture

Immortalized fallopian tube hTERT-FT and MeT-5A cells were purchased from ATCC, and the lentiviral transduction method was followed to express the red fluorescent protein (RFP)^48^. The hTERT-FT cells were cultured in DMEM F12 media (HiMedia, AT007F), added with 10% fetal bovine serum. MeT-5A cells were cultured in Medium 199 media (HiMedia, AL057A) along with 10% fetal bovine serum (Gibco; 10270), hydrocortisone, sodium selenite, and insulin. The cells were maintained in a 5% carbon dioxide, 37° C temperature, humidified incubator.

### Animal experiments

Female BALB/c mice aged 4–6 weeks were used, with all procedures conducted in compliance with ethical guidelines and institutional approval (CAF/Ethics/106/2024).

### Hydra culturing

*Hydra magnipapillata* (Gift from Chinmoy Patra, Aghakar Research Institute, Pune) were cultured in hydra medium (1 mM NaCl, 1 mM CaCl_2_, 0.1 mM KCl, 0.1 mM MgSO_4_, 1 mM Tris, pH 7.6) at 18 °C. Hydra were fed freshly hatched Artemia nauplii every day and cleaned 4-5 hours after feeding.

### Determination of LD_50_ values of MGO and TA

The inhibition concentration 50 (LD_50_) was estimated using different concentrations of MGO and TA. A 10 mM stock solution of MGO and TA was prepared using hydra medium. For MGO, test concentrations ranged from 100 to 2400 µM, and for TA from 5-60 µM, freshly prepared every day using hydra medium. Cadmium chloride (3 µM) was used as the positive control, while hydra medium was used as the negative control. All hydra were starved for 48 hours before the experiment. Ten hydras were immersed in 1mL of the desired concentration of each test solution in a 24-well plate. Each concentration was tested in three biological replicates. The experiment was carried out over 4 days at 18 °C in the dark. The solutions were replaced with freshly made solutions at the same concentrations every 24 hours. All the hydra were microscopically observed, and the number of live hydras was recorded every 24 hours. Absence of response to stimulus or dissociation of cells indicated death.

### Preparation of Collagen I and Laminin-rich Basement membrane (LrBM) gels

Collagen I gels were prepared from the previously reported method^16^. Briefly, 330 µL of Collagen I (3 mg/mL) was added to the 560 µL of 1X PBS, and 100 µL of 10X DMEM medium (HiMedia AT006F). The mixtures were properly mixed, followed by neutralization with 6 µL of 2N NaOH. 50 µL of prepared collagen solution was added to the 24-well plate. The gels were polymerized at 37° C for 60 minutes, followed by an overnight 4° C incubation to further enhance the polymerization. Matrigel (50 µL of 9.3 mg/mL stock concentration) was added as spherical beads into the 24-well plate. The gels were incubated at 37° C for 60 minutes and used for further studies.

### Picrosirius staining of MGO and TA-treated Collagen I gels

The prepared collagen gels were incubated with control (1X PBS), 100 µM of TA (1 day), 200 µM of MGO (4 days), and 100 µM of TA (1 day) followed by 200 µM MGO (4 days). After 4 days of incubation, the solution was removed and thoroughly washed with 1X PBS. The picric acid solution was prepared as per the previous report and was added to the gels and incubated for 2 hours, followed by washing with 0.05% acetic acid and 100% ethanol^22^. Finally, the gels were washed with 1X PBS to remove the excess stain.

### Crystal violet staining of MGO and TA-treated LrBM gels

The prepared LrBM gels were incubated with control (1X PBS), 100 µM of TA (1 day), 200 µM of MGO (4 days), and 100 µM of TA (1 day) followed by 200 µM MGO (4 days). After the 4 days of incubation, the solution was removed and thoroughly washed with 1X PBS, followed by formaldehyde fixation (1%) at 37° C for 60 minutes. Formaldehyde solution was removed and thoroughly washed with 1X PBS. The crystal violet was added to the LrBM and incubated for 30 minutes, the stain was removed, and twice washed with 1X PBS.

### Cell adhesion assay MGO and TA-treated Collagen I gels and LrBM gels

For the cell adhesion assay, collagen gels were prepared on the chamber well plates. Briefly, 100 µL of neutralized collagen was coated on the well and polymerized at 37° C for 60 minutes, followed by an overnight 4° C incubation. Similarly, 100 µL of Matrigel was coated on the well and polymerized at 37° C for 60 minutes. Both the gels were incubated with control (1X PBS), 100 µM of TA (1 day), 200 µM of MGO (4 days), and 100 µM of TA (1 day) followed by 200 µM MGO (4 days). The solution was removed and washed thrice with 1X PBS. The RFP labelled hTERT-FT and MeT-5A cultured cells were added to each well (10000 cells per well). The cells were maintained in a 5% carbon dioxide, 37° C temperature, humidified incubator for 45 minutes. The cell suspension was removed and washed twice with 1X PBS, followed by glutaraldehyde fixing (0.02%) at 37° C for 30 minutes. The aldehyde solution was removed and washed thrice with 1X PBS, and cells were stored at 4° C for further use. The Olympus IX83 inverted fluorescence microscope (10X magnification) was used for imaging of adhered cells, and ImageJ software was used for processing and cell counting analysis.

### Cell spreading assay on MGO and TA-treated Collagen I and LrBM gels

For the cell spreading assay, collagen gels were prepared on the chamber well plates. Briefly, 100 µL of neutralized collagen and matrigel were coated on the well and polymerized at 37° C for 60 minutes. The gels were incubated with control (1X PBS), 100 µM of TA (1 day), 200 µM of MGO (4 days), and 100 µM of TA (1 day) followed by 200 µM MGO (4 days). The solution was removed and washed thrice with 1X PBS. Immortalized hTERT-FT and MeT-5A cultured cells were added to each well (10000 cells per well). The cells were maintained in a 5% carbon dioxide, 37° C temperature, humidified incubator for 6 h. The media was removed and washed twice with 1X PBS, followed by formaldehyde fixing (4%) at 37° C for 30 minutes. The formaldehyde solution was removed and washed three times with 1X PBS. The fixed cells were stained using DAPI (10 minutes) and Phalloidin (12 h) for DNA and F-actin, respectively. The Zeiss microscope (20X magnification) was used for imaging of cells, and ImageJ software was used for image processing.

### Preparation of the cell-extracted extracellular matrix (CEEM) by cell lysis

The ammonium hydroxide cell lysis method was followed for the preparation of the extracellular matrix. Immortalized hTERT-FT cells (200000 cells per well) were cultured till confluency in the 6-well plate and allowed to grow for 72 hours in a 5% carbon dioxide, 37° C temperature, humidified incubator. After 72 h, the media was removed, and cells were washed twice with 1X PBS. Then, 20 mM of ammonium hydroxide solution (25%) was added to each well and allowed to incubate for 5 minutes. The solution was removed and thoroughly washed with autoclaved filtered distilled water and stored at 4° C for further use.

### Crystal violet staining of MGO and TA-treated CEEM

The above prepared CEEM was incubated with control (1X PBS), 100 µM of TA (1 day), 200 µM of MGO (4 days), and 100 µM of TA (1 day) followed by 200 µM MGO (4 days). After the 4 days of incubation, the solution was removed and thoroughly washed with 1X PBS, followed by formaldehyde fixation (1%) at 37° C for 60 minutes. Formaldehyde solution was removed and thoroughly washed with 1X PBS. The crystal violet was added to the plate and incubated for 30 minutes; the stain was removed and twice washed with 1X PBS. The Olympus IX83 inverted fluorescence microscope (10X magnification) was used for imaging of CEEM.

### Cell adhesion assay on MGO and TA-treated CEEM

The CEEM was incubated with control (1X PBS), 100 µM of TA (1 day), 200 µM of MGO (4 days), and 100 µM of TA (1 day), followed by 200 µM MGO (4 days). The solution was removed and washed thrice with 1X PBS. The RFP labelled FT and MeT-5A cells were added to each well (10000 cells per well). The cells were maintained in a 5% carbon dioxide, 37° C temperature, humidified incubator for 45 minutes. The cell suspension was removed and washed twice with 1X PBS, followed by glutaraldehyde fixing (0.02%) at 37° C for 30 minutes. The glutaraldehyde solution was removed and washed thrice with 1X PBS. The Olympus IX83 inverted fluorescence microscope (10X magnification) was used for imaging of adhered cells, and ImageJ software was used for processing and cell counting analysis.

### Cell spreading assay on MGO and TA-treated CEEM

For the cell spreading assay, the CEEM was incubated with control (1X PBS), 100 µM of TA (1 day), 200 µM of MGO (4 days), and 100 µM of TA (1 day) followed by 200 µM MGO (4 days). The solution was removed and washed thrice with 1X PBS. Immortalized hTERT-FT and MeT-5A cultured cells were added to each well (10000 cells per well). The cells were maintained in a 5% carbon dioxide, 37° C temperature, humidified incubator for 6 h. The media was removed and washed twice with 1X PBS, followed by formaldehyde fixing (4%) at 37° C for 30 minutes. The formaldehyde solution was removed and washed three times with 1X PBS. The fixed cells were stained using DAPI (10 minutes) and Phalloidin (12 h) for DNA and F-actin, respectively. The Olympus IX83 inverted fluorescence microscope (10X magnification) was used for imaging of adhered cells, and ImageJ software was used for processing and cell counting analysis.

### Preparation of cell lysate of MGO and TA-treated cells

The RIPA/cell extraction buffer was used to lyse the cells. Immortalized hTERT-FT cells (200000 cells per well) were seeded onto the 6-well plate. Cells were treated with media (control), 300 µM of MGO was added along with media for 48 hours, 10 µM of TA was added along with media for 24 hours, and 10 µM of TA was added along with media for 24 hours, followed by 300 µM MGO added along with media for 48 hours. The cells were maintained in a 5% carbon dioxide, 37° C temperature, humidified incubator. After the treatment media was removed and the cells were washed with ice-cold 1X PBS twice. The well plate was kept on ice, and 800 mL of RIPA/cell extraction buffer was added to the plate. Cells were scraped using a cell stainer, and the solution was transferred to centrifuge tubes and gently vortexed every 10-minute intervals for 40 minutes. The lysate solution was centrifuged at 12000 rpm for 20 minutes at 4° C, and the supernatant solution was collected and stored at −20° C for further use.

### Biochemical characterization of cell lysate

#### Estimation of protein carbonyl group

Glycated and TA-pretreated samples (500 µL) were mixed with 500 µL of 20% trichloroacetic acid and 500 µL of 10 mM 2,4-dinitrophenyl hydrazine prepared in 2.5 M hydrochloric acid, which specifically reacts with carbonyl groups to form a colored hydrazone complex^49^. The mixture was incubated at room temperature for 60 minutes, followed by centrifugation at 7000 rpm for 5 minutes. The pellet was washed with 500 µL of ethanol and an ethyl acetate mixture (1:1). 1 mL of 6M urea was used to resuspend the pellets, and the solution was centrifuged at 7000 rpm for 10 minutes. The absorbance was measured at 365 nm.

#### Estimation of the free amino group

For the estimation of the amino group, 100 µL of the lysis sample was mixed with 100 µL of phosphate buffer (pH 7.2) and 100 µL of 0.1 M para-benzoquinone dissolved in DMSO^50^. The mixture was incubated at room temperature for 5 minutes, and absorbance was measured at 480 nm.

### Biochemical characteristics of MGO and TA-treated CEEM

The prepared extracellular matrix was incubated with control (1X PBS), 100 µM of TA (1 day), 200 µM of MGO (4 days), and 100 µM of TA (1 day) followed by 200 µM MGO (4 days). The solution was removed and washed thrice with 1X PBS, followed by formaldehyde fixation (1%) at 37° C for 60 minutes. Formaldehyde solution was removed and thoroughly washed with 1X PBS. The samples were treated with crystal violet for 90 minutes, then the stain was removed and twice washed with 1X PBS, followed by a methanol wash. The protein concentration of the samples was calculated by measuring absorbance at 595 nm. The carbonyl group and free amino group of the protein were estimated by above mentioned method.

### *Ex vivo* studies

For the *ex vivo* experiments, mesenteric tissues were dissected from the 4-6-week-old BALB/c female mice. Mice were sacrificed by cervical dislocation. The mesenteries were surgically removed from the abdomen and strung using surgical thread to the transwell. The transwells were then placed in a 24-well plate containing media, 200 µM of MGO in media (2 days), 10 µM of TA in media (1 day), and 10 µM of TA in media (1 day), followed by 200 µM MGO in media (2 days). The mesenteries were maintained in a 5% carbon dioxide, 37° C temperature, humidified incubator. After the treatment media was removed and washed twice with 1X PBS, followed by formaldehyde fixing (4%) at 4° C for 60 minutes. The formaldehyde solution was removed and washed three times with 1X PBS. The mesenteries were permeabilized with 0.5 % Triton-X in PBS for 5 h, followed by blocking with 0.3 % BSA, 0.1% Triton-X in 1X PBS at 4° C for 2 h. The mesenteries were stained using DAPI (10 minutes) and Phalloidin (12 h) for DNA and F-actin, respectively. The Zeiss microscope (10X magnification) was used for imaging of mesenteries, and ImageJ software was used for image processing.

### Statistics

The GraphPad Prism 10 was used for data analysis and plotting graphs. Significance computed using one-way ANOVA with Tukey’s post hoc pairwise comparisons.

### Instrumentation

#### Circular Dichroism (CD) studies

For CD analysis, pre-polymerized collagen (pH 4.7, 0.1 mg/mL) was treated with 200 µM MGO, 50 µM TA, and 200 µM MGO-50 µM TA. The solutions were kept for overnight incubation at 4° C. The analysis was performed on a Jasco-815 spectropolarimeter purged with inert gas in the spectral range of 190–260 nm in a rectangular quartz cuvette with an optical path of 0.1 cm.

### NMR sample preparation

For the titration experiments, 18 different samples of 1 mL were prepared in a 10 mM phosphate buffer, pH 7.4. For all the samples, the MGO concentration was 200 μM. The TA concentrations were 0, 4, 8, 16, 24, 32, 40, 56, 64, 72, 80, 120, 160, 200, 400, 600, 800, and 1000 μM. The samples were incubated for 24 hours and freeze-dried for 4 hours. NMR samples were made by adding 588 μL of D_2_O and 12 μL of 100 μM 3-(trimethylsilyl) propionic acid (internal chemical shift and intensity standard) in D_2_O to the freeze-dried samples. As the 1 mL samples were freeze-dried and subsequently re-dissolved in 0.6 mL of D₂O for NMR analysis, the absolute concentrations of all components in the NMR samples increased 1.6-fold, while their molar ratios remained unchanged.

### NMR spectroscopy and data analysis

NMR titration experiments were acquired at 300 K using a Bruker Avance Neo 14.1 T (600 MHz ^1^H Larmor frequency) spectrometer equipped with a triple resonance broadband probe (iTBO probe). The ^1^H 1D spectrum of MGO was recorded on a Bruker Avance Neo 18.8 T (800 MHz ^1^H Larmor frequency) spectrometer equipped with a TXI probe at 298 K. All ^1^H 1D experiments were recorded by averaging over 128 transients using 25k points over a spectral width of 12 kHz and a relaxation delay of 2s. The residual water was suppressed through an excitation sculpting scheme.

^1^H 1D spectra were processed and analysed using the Topspin 4.1.3 software package. All FIDs were zero-filled to 65k prior to the Fourier transformation. The chemical shift of the internal standard was referenced to 0 ppm, and its peak intensity was normalized to 100. The titration plot was generated by plotting the intensity of the analyte peak relative to the internal standard (relative intensity) as a function of the ratio of methylglyoxal and tannic acid concentrations ([TA]/[MGO]). The noise in the spectrum was used as an error in intensity and estimated from the peak intensity of the internal standard and its signal-to-noise ratio (SNR). The SNR values were obtained using the SINO macro in Topspin, for which 0.01 ppm around the internal standard peak was used as the signal region and 9.6 ppm to 10 ppm was used as the noise region.

### Interaction model and NMR peak intensity analysis

The titration plots were fitted to an n-site binding model where each molecule of TA can associate with n molecules of MGO. The reaction is given by the equation,

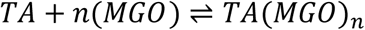

The association constant (K_a_) is defined as follows.

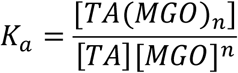

Since the three hydrated forms of MGO are in equilibrium, it is assumed that the peak intensities of the three MGO forms respond similarly to the TA titrations. The titration profiles of different hydrated forms of MGO were fit globally to this n-site binding model using an in-house Python script that numerically solves the equilibrium and mass-balance equations to derive the intensities of unbound MGO at each titration point.

## Author Contributions

R.B. conceived the project and designed the experiments. C.I. performed the experiments. NMR experimental analysis and data curation were done by C.N.V. and A.S. Hydra experimental analysis and data curation were done by P.A. and S.U. R.B. and C.I. wrote the paper with input from all authors.

## Data Sharing

Raw data generated for the paper will be shared upon reasonable request.

